# Target of Rapamycin drives unequal responses to essential amino acid depletion in egg laying

**DOI:** 10.1101/2021.11.08.467165

**Authors:** André N. Alves, Carla M. Sgrò, Matthew D. Piper, Christen K. Mirth

**Affiliations:** School of Biological Sciences, Monash University, Melbourne, VIC 3800, Australia

**Keywords:** Cellular signalling, *Drosophila melanogaster*, egg laying dynamics, Target of Rapamycin (TOR), GCN2, egg chamber development

## Abstract

Nutrition shapes a broad range of life history traits, ultimately impacting animal fitness. A key fitness-related trait, female fecundity is well known to change as a function of diet. In particular, the availability of dietary protein is one of the main drivers of egg production, and in the absence of essential amino acids egg laying declines. However, it is unclear whether all essential amino acids have the same impact on phenotypes like fecundity. Using a holidic diet, we fed adult female *D. melanogaster* diets that contain all necessary nutrients except one of the 10 essential amino acids and assessed the effects on egg production. For most essential amino acids, depleting a single amino acid induced as rapid a decline in egg production as when there were no amino acids in the diet. However, when either methionine or histidine were excluded from the diet, egg production declined more slowly. Next, we tested whether GCN2 and TOR were involved in this difference in response across amino acids. While mutations in GCN2 did not eliminate the differences in the rates of decline in egg laying among amino acid drop-out diets, we found that inhibiting TOR signalling caused egg laying to decline rapidly for all drop-out diets. TOR signalling does this by regulating the yolk-forming stages of egg chamber development. Our results suggest that amino acids differ in their ability to induce signalling via the TOR pathway. This is important because if phenotypes differ in sensitivity to individual amino acids, this generates the potential for mismatches between the output of a pathway and the animal’s true nutritional status.

## Introduction

Diet impacts most life history traits in animals. It determines their ability to survive, resist stress, grow, and even dictates how many offspring they will have (Simpson and Raubenheimer, 2012). The term diet typically refers to a combination of nutrients, each of which have distinct impacts. For example, a high protein diet can reduce lifespan by negatively affecting gut function, whereas a high carbohydrate, low protein diet prolongs lifespan by reducing egg production (Zanco et al., 2021; Regan et al., 2016). How diet alters these life history traits has been difficult to pin down, because different components of the diet are sensed by distinct signalling pathways and because the effects of diet differ across organs. However, it is important for animals to sense the components of the diet accurately, so that they can adjust their phenotypes in a manner appropriate to the nutrients available. Failure to do so would lead to a mismatch between the phenotype and the diet.

The cells within organs respond to nutrients via nutrient-sensing signalling pathways. Signalling pathways that respond to macronutrients have been identified, including those that respond to protein concentrations. Cells directly sense amino acid concentration through the General Control Non-derepressible 2 (GCN2) and Target of Rapamycin (TOR) pathways. GCN2 is a ubiquitous protein kinase activated in the presence of uncharged tRNAs, which occurs when there is an insufficiency of amino acids (Armstrong et al., 2014; Sonenberg and Hinnebusch, 2009). In these conditions, GCN2 becomes active and signals to inhibit general protein translation (Sonenberg and Hinnebusch, 2009). The TOR pathway also responds to the intracellular concentration of amino acids, increasing cell survival, growth, and proliferation in the presence of amino acids (Wang and Proud, 2009). While there is some suggestion in the literature that GCN2 and TOR might not respond with the same sensitivity to all amino acids, most of these differences have only been characterised in cell culture (Yuan et al., 2017; Ye et al., 2015).

One way to uncover differences in sensitivity in the GCN2 and TOR pathways to individual amino acids is to examine how life history traits that are easy to quantify respond to deficits in individual essential amino acids. Female fecundity is an excellent choice for such studies. The effect of protein on female fecundity has been extensively studied (Mirth et al., 2019; Lee et al., 2008; Piper et al., 2014; Armstrong et al., 2014). This trait is convenient because we can trace all the effects of dietary protein to a single output, the number of eggs laid or offspring produced. The effects of protein concentration on fecundity are well documented, consistent, and transferable across different classes of animals. For example, in several species of fish, such as *Oreochromis niloticus* and *Poecilia reticulata*, the concentration of dietary protein in females is positively correlated with the number of offspring generated (Dahlgren, 1980; Hafedh et al., 1999). This is also true for insects, where reducing the concentration of protein in the food decreases egg production in various species, including carabid beetles, locusts, crickets, Queensland fruit-flies, and *Drosophila* fruit flies (Wallin et al., 1992, Behmer et al., 2001, Maklakov et al., 2008, Lee et al., 2008, Fanson & Taylor, 2011). Indeed, protein is often a limiting nutrient for fecundity across the animal kingdom (White, 1993).

Studies in the fruit fly, *Drosophila melanogaster*, have contributed significantly to what we know about how protein exerts its effects on egg laying. The rate of egg laying is highest when flies ingest protein-rich food (Lee et al., 2008; Grandison et al., 2009; Piper et al., 2014). Thus, protein availability limits fecundity in *D. melanogaster* like it does in other animals. *D. melanogaster* bear the additional advantages of a fast life cycle, a range of genetic tools, a fully synthetic diet (Piper et al., 2014), and extensive studies on the development of eggs, making it possible to test whether fecundity differs in sensitivity across amino acids and to identify the signalling pathway responsible for these differences.

*D. melanogaster* require twenty amino acids to make up all proteins encoded in the genome. Of these, there are ten essential and ten non-essential amino acids. An essential amino acid has to be obtained from the diet, since it cannot be synthesised by the organism (Wu et al., 2013). In the wild, *D. melanogaster* must obtain essential amino acids by eating yeast proliferating in rotting fruit. The absence of any one of the ten essential amino acids stops egg development (Sang and King, 1961), and females stop laying eggs (Grandison et al., 2009).

Egg development in *D. melanogaster* occurs within the ovaries but depends on yolk protein and other signals from the fat body and hormones from the brain and *corpora allata* (Armstrong et al., 2014). *D. melanogaster* ovaries are made up of 16-22 ovarioles (David, 1970; Hodin and Riddiford, 2000; Mendes and Mirth, 2016; Sarikaya et al., 2011), strings of developing eggs and their surrounding somatic cells. At the anterior tip of each ovariole, the terminal filament cells, cap cells, and escort cells compose the stem cell niche for 2-3 germline stem cells (Spradling, 1993). These germline stem cells adhere to the supporting cap and escort cells, which play an important role in ensuring the stem cells remain undifferentiated (Koch and King, 1966). As the germline stem cells divide, the daughter cell that remains attached to the cap cells remains a stem cell, while the other daughter cell develops into a cystoblast (Koch and King, 1966). The cystoblast will divide 4 times without completing cytokinesis, resulting in 16 cells that share cytoplasmic connections called germline cysts (Koch and King, 1966).

As the cyst moves out of the germarium, it becomes encapsulated by a single layer of follicle cells, which are produced from divisions of the follicle stem cells (Koch and King, 1966). Once encapsulated, the cysts are known as egg chambers. Egg chambers go through 14 stages of development to become fully formed oocytes (King et al., 1956; Cummings and King, 1969, Figure 1). The first seven stages can be broadly grouped as the early stages of egg chamber development. These early stages are characterised by growth of the egg chamber, and differentiation of one of the sixteen germline cells into an oocyte while the remaining cells take on the role of nurse cells (Brown and King, 1964). Stages 8-11 are the vitellogenic, or yolk-forming, stages. During the yolk-forming stages the oocyte fills up with yolk produced by the fat body and follicle cells. The nurse cells begin to deposit protein and mRNA into the oocyte that will be necessary for embryogenesis (Cummings and King, 1969). The late stages of egg chamber development involve depositing the egg membranes, the death of the follicle and nurse cells, and building the dorsal filaments, which are tubes that protrude from the anterior of the egg (Cummings and King, 1969). Early, yolk-forming, and late stages of egg development all depend on dietary protein to progress (Drummond-Barbosa and Spradling, 2001).

**Figure 1:**
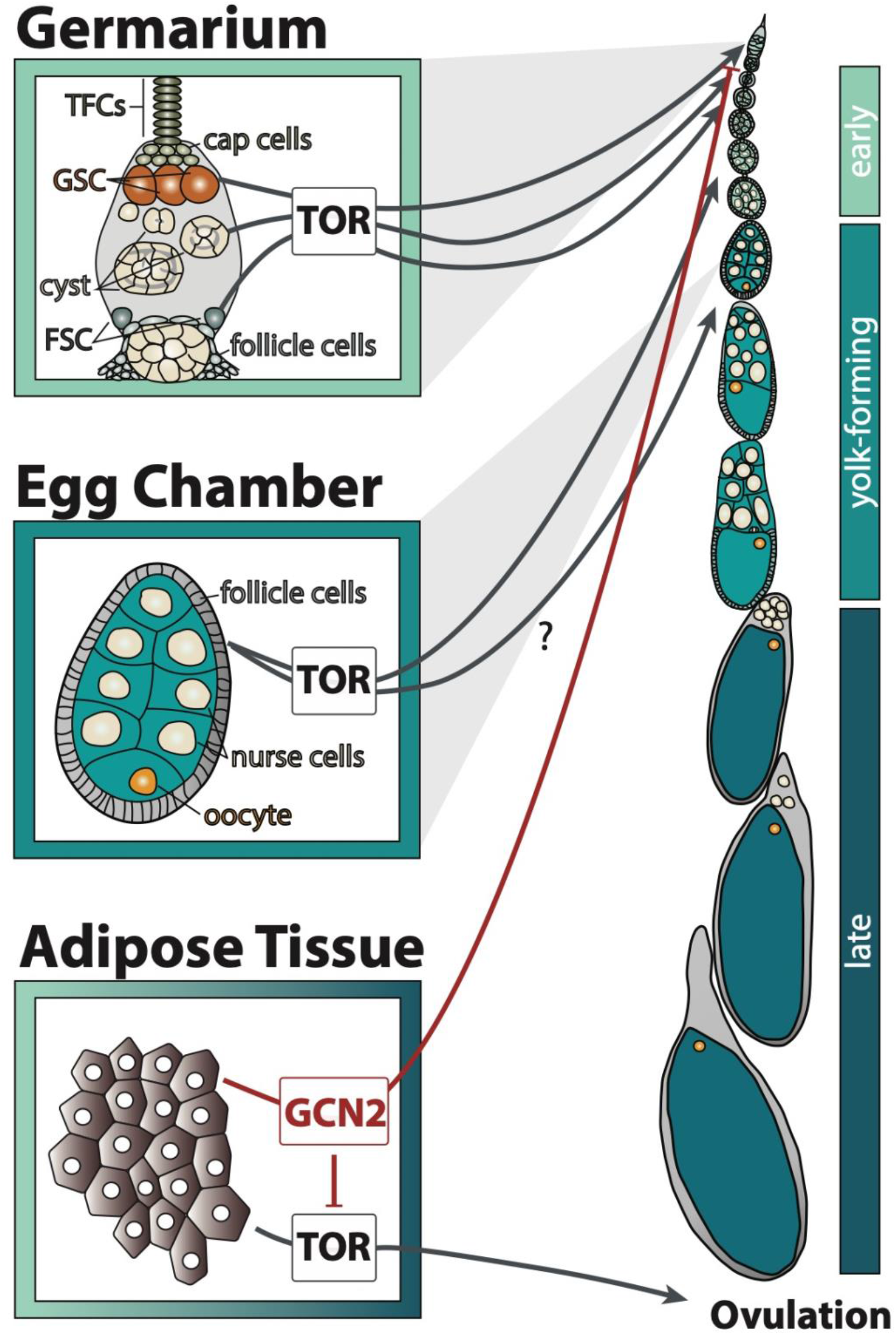
GCN2 and TOR signalling in the adipose tissue and ovary transmit amino acid availability by affecting distinct processes of egg chamber development. Early stages include stages 1-7, yolk-forming includes stages 8-11, and late includes stages 12-14 of egg chamber development. The question mark indicates that it is unclear through which cell type TOR regulates the yolk-forming stages. TFCs: terminal filament cells, GSC: germline stem cells, FSC: follicle stem cells.

Like other cells in the body, GCN2 and TOR signalling regulates egg chamber development in response to amino acid availability. TOR is required in many cell types of the ovary, and also in the fat body, to promote egg development. When intracellular amino acids are present at high concentrations, TOR is active in the germline stem cells to promote the maintenance of stem identity and to induce their division (LaFever et al., 2010, Figure 1). Before the cyst leaves the germarium, TOR is important for cyst growth (LaFever et al., 2010). It also promotes follicle stem cell division and follicle growth, necessary for egg chamber growth (LaFever et al., 2010). The egg chambers of TOR mutant animals never initiate yolk formation. It is unclear if this is due to TOR activity in the follicle cells, fat body, or in other tissues (LaFever et al., 2010).

Amino acid sensing in the fat body further regulates germline stem cell maintenance and ovulation (Armstrong et al.,2014). The fat body regulates germline stem cell number in response to low amino acids via GCN2, but not TOR (Armstrong et al.,2014, Figure 1). Low amino acid concentrations activate GCN2 in the fat body, which in turn represses TOR and blocks ovulation (Armstrong et al.,2014, Figure 1). The signal secreted by the fat body to control these processes in the ovary is unknown. Nevertheless, because GCN2 and TOR perform distinct roles in regulating egg chamber development (Figure 1), we can use these differences to discern if these two pathways differ in sensitivity to individual amino acids.

To investigate differences in the sensitivity of egg production across essential amino acids, we used a fully synthetic diet that allows us to eliminate individual amino acids and determine their specific impact on egg laying (Piper et al., 2014). We hypothesized that if amino acids differed in their ability to trigger cellular signalling, then eliminating individual amino acids could induce different rates of decline in egg laying. If egg laying did decline at different rates, we could then probe whether this resulted from the activity of either the GCN2 or TOR signalling and define the precise effects on egg development. Our work highlights that these key nutrient signalling pathways are limited in their ability to faithfully represent amino acid availability, which has the potential to lead to mismatches between diet and phenotype.

## Methods

### Stocks and Fly Maintenance

We used three outbred populations of flies in this study: a population of wild-type flies (Red Dahomey), an outbred Dahomey population that also carries a mutation in the white gene (w^1118^, hereafter referred to as white Dahomey - wDah), and a white Dahomey population that harbours a null mutation in GCN2 (GCN2Δ, supplied from Linda Partridge and Sebastian Grönke). Both Red Dahomey and White Dahomey were maintained in population cages with overlapping generations at 25ºC on a 12h light:dark cycle. GCN2Δ were maintained in vials at 18ºC on a 12h light:dark cycle. All populations were maintained on sugar, yeast, agar food (SYA) as described in Bass et al., 2007.

### Staging and collecting adults

To control for larval rearing density and to synchronize adult emergence time, we collected eggs from the parental generation by leaving them to lay in embryo collection cages (Genesee Scientific) on 90mm petri dishes half-filled with apple juice/agar medium, as described in Linford et al., 2013, for 24 h at 25ºC. 250 eggs were then distributed into each food bottle, which contained SYA food at 25ºC.

The adult flies that emerged from these cultures were collected over a 48-hour period, transferred to new bottles containing SYA medium, and left to mate for 48 h. Once mated, five female flies were transferred into vials that contained a fully synthetic diet (100N, named for having 100 mM of biologically available nitrogen) (Piper et al. 2017), grouped in ten replicates using drosoflippers (http://drosoflipper.com/), and left to acclimatize for one week, replacing the vials with fresh food every three days. Synthetic diets were made according to Piper et al. 2014.

### Diets, drug treatments, and egg counts

After a week of acclimatization to the synthetic diet, females were transferred to new vials that contained one of twelve diets. In addition to two control diets - all amino acid diet (All AA) and no amino acid diet (No AA) - we used ten experimental diets that contained all but one of ten essential amino acids: arginine^-^, histidine^-^, isoleucine^-^, leucine^-^, lysine^-^, methionine^-^, phenylalanine^-^, threonine^-^, tryptophan^-^, and valine^-^. We established ten replicate vials per diet. Females were transferred to fresh diet every 24 h, after which the eggs laid over the previous 24 h period were counted manually. This was done for 7 to 9 consecutive days.

Mutations in genes from amino acid sensing signalling pathways are known to reduce ovariole number, thereby limiting the maximum number of eggs a female can produce (Green & Extavour, 2014). To account for differences in ovariole number between the GCN2Δ and control genotypes (wDah), the ovaries of 10 to 15 females of each genotype were dissected out in cold PBS and their ovarioles were counted.

To inhibit the mTOR pathway, Rapamycin was diluted in pure ethanol to make a 900 μM stock solution. For each diet, 100 μL of the solution was dispensed on the surface of 3mL of food to a final concentration of 30 μM. The same quantity of the carrier, pure ethanol, was added to a control set of diets.

### Ovary Dissection, Imaging, and Staging

For ovary dissections, flies were collected as described above, and after a week of acclimatization on an all amino acid diet (All AA) additional groups of females from the three genotypes mentioned above were transferred to new vials that contained one of six diets. In addition to two control diets – complete amino acid diet (All AA) and no amino acid diet (No AA) – we used four experimental diets that contained all but one of four essential amino acids: arginine^-^, histidine^-^, methionine^-^ and phenylalanine^-^. Females were transferred to fresh diet every 24 h. At days zero, three, and seven, five replicates of five flies were removed and their abdomens dissected. Red Dahomey flies were divided between Rapamycin- and ethanol-laced diets, as described above.

We dissected the abdomens in cold phosphate-buffered saline (PBS). Female abdomens were then fixed in 4% paraformaldehyde (Sigma) overnight at 4ºC. After fixing, abdomens were washed with phosphate-buffered saline with 0.1% Tween (PBST) four times for 15 minutes before RNase (Promega, original concentration of 4 mg/mL) was added in a concentration of 10 μL/mL for 20 minutes. The samples were then washed four times before adding DAPI at a concentration of 0.1 μL/mL for 5 minutes. After staining, two final washes with PBST were performed, followed by two extra washes with PBS. Following staining, ovaries were dissected from abdomens and each treatment placed in one well of a 96-well plate with Fluoromount-G.

Imaging was done through a Leica SP5 5-Channel, using a resonant scanner, with a 20x objective. File conversion from Leica Matrix Screener was done through Python app LM2BS (https://github.com/VolkerH/LeicaMatrixScreener2BigStitcher). H5 files were then read and stitched through the BigStitcher plug-in available in FIJI (version 2.0) and saved as TIFF files.

Ovaries were staged based on characteristics described in (Jia et al., 2016). Only ovaries with clearly visible ovarioles through the z-axis were considered adequate to stage. Staging was grouped into one of three categories: early stages (stages 1-7), yolk-forming stages (stages 8-11), and late stages (stages 12-14).

### Data Analysis

To test for differences in total eggs laid between diets and genotypes, we fit a generalised linear model with the total eggs laid per female as a response variable, with diet and genotype/treatment (where appropriate) as fixed effects, assuming a Poisson distribution with a log link function. Post-hoc comparisons of the means were conducted using the emmeans function (emmeans package).

We identified the models that best fit the effect of amino acid drop-out diets on the number of eggs laid over time by initially fitting the data on the 0N diet using either a linear model, second- or third-degree polynomial models, or a self-starting logistical (SSlogis, *y* = *asym*/(1 + *e*^((*xmid−day*)/*scal*)^) regression model using the nls package, where asym is the asymptote, xmid is the flexion or mid-point of the curve, and scal is the scaling coefficient (Bates and Watts, 1988, Chambers and Bates, 1992). AIC and BIC were calculated to assess which model best fit the data.

To compare the dynamics of the curve across diets, we used logistic regression (SSlogis), which provides easy to interpret parameters that correspond to rate of change (scaling coefficient) and midpoints (xmid coefficient) of the data. We grouped diets into two groups: the first group where amino acid drop-out diet induced a decline in egg production at a similar rate as the 0N, and a second group where egg production declined more slowly. We then tested whether fitting the logistic regression with coefficients specific to each group improved the model fit over a null model fit with common coefficients. We tested model fit using partial F tests, AIC, and BIC.

Because it is difficult to assess interactions between diets and genotypes/treatments with logistic regression models, we analysed interactions between diet, time (day), and genotype/treatment using third order polynomial regressions. To do this, we fit the number of eggs laid per female as a response variable and a third polynomial fixed effect of day interacting with diet. Where appropriate, genotype or treatment was included as a fixed effect. Replicate was used as a random effect within each day. Emtrends was used to obtain statistical differences between slopes of each diet.

To compare egg chamber categories and respective percentages, we fit the data with generalised linear models, assuming a binomial distribution. The proportion of egg chambers at a particular stage was used as a response variable, and day, category, diet and genotype/treatment (where appropriate) were used as fixed effects. We conducted post hoc comparisons of means and slopes using emmeans and emtrends (emmeans package), respectively.

Data was analysed and visualized in R Studio (version 3.4.1). Plots were produced using ggplot2 (tidyverse package, Wickham et al., 2019). All data and scripts are available in Figshare (reference number to be provided).

## Results

### The number of eggs laid differs between amino acid drop-out diets

Based on previous data (Piper et al., 2014), we hypothesised that the elimination of individual essential amino acids will create different rates of decline in egg laying. To test this, we maintained five female flies on each of twelve different synthetics diets and assessed the effect on the number of eggs laid during nine days of exposure to the treatment diets. These diets included a diet containing all essential and non-essential amino acids (All AA), a negative control diet with no amino acids (No AA), and ten different drop-out diets containing all except one of the essential amino acids.

We observed that in the absence of all amino acids (No AA), egg laying dropped sharply to zero by the fourth day (Figure 2A). In contrast, female flies fed on diets containingthe full complement of amino acids (All AA) showed a gradual decline of ~8% per day in egg laying. The single amino acid drop-out diets induced one of two different responses in egg laying. For seven of these diets, including the arginine, isoleucine, leucine, lysine, phenylalanine, threonine, and tryptophan drop-out diets, egg laying declined at a similar rate as it did on the diet without any amino acids (Figure 2A). As a result, over the nine days examined these females laid similar numbers of eggs as those on the diet without amino acids (Figure 1B). However, for the histidine, methionine, and valine drop-out diets, egg laying declined more gradually with time (Figure 2A), resulting in significantly higher numbers of eggs laid over the nine-day period (Figure 2B, GLM: χ^2^_114,119_ = 175, p < 0.001).

**Figure 2:**
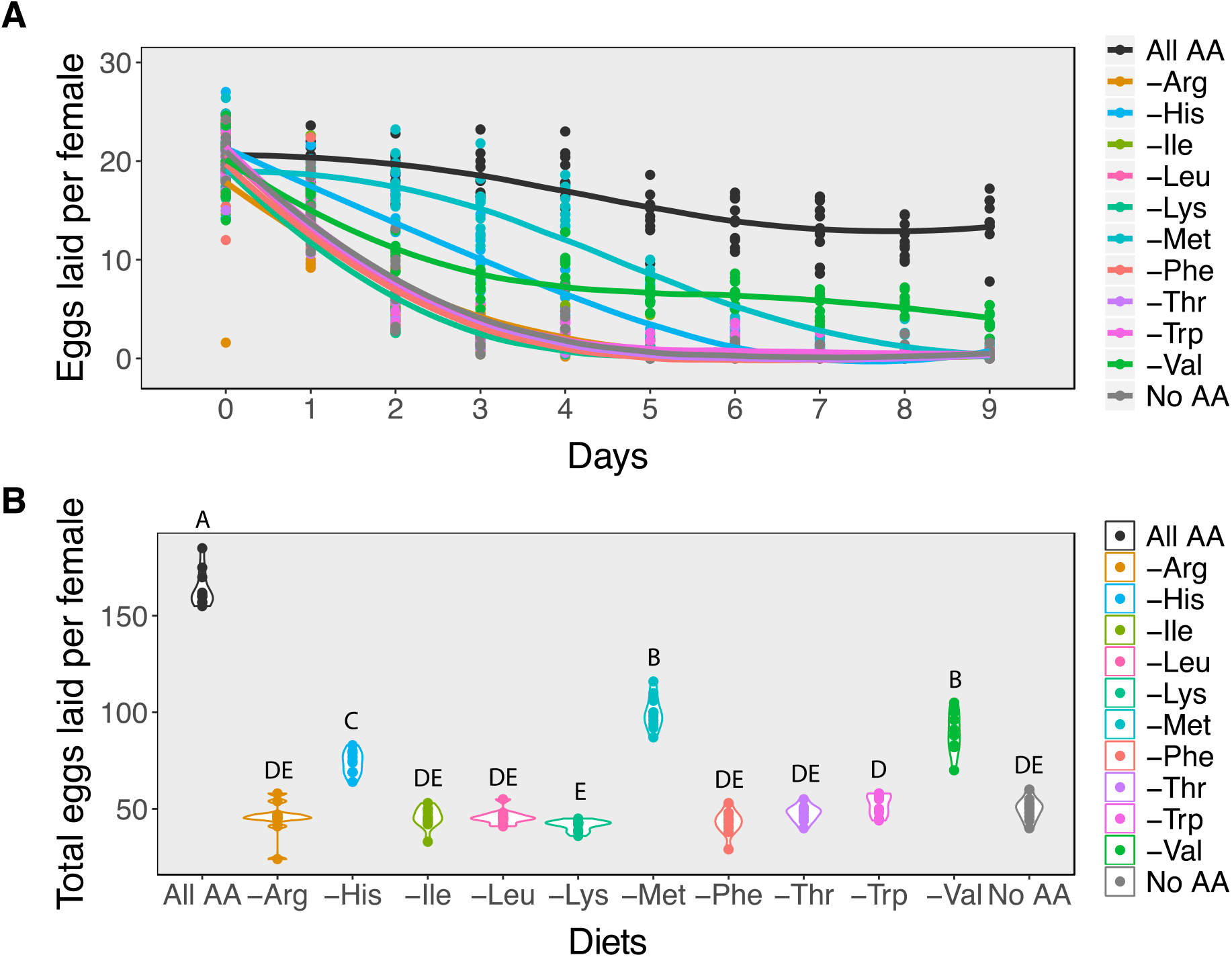
Amino acids differ in their effects on egg laying. A) While in most cases removing individual essential amino acids results in a rapid decline in egg laying at a rate similar to removing all amino acids from the diet, removing valine, histidine, or methionine results in a slower rate of decline. B) For most diets in which a single amino acid is removed, the total number of eggs that females produce over a nine-day period is similar to females on a diet that contains no amino acids. Females on either histidine, valine, or methionine drop-out diets produce more eggs than those on diets missing any of the other essential amino acids. Diets that have differing letters are significantly different, as determined from post hoc pairwise comparisons derived from a generalised linear model assuming a Poisson distribution.

To test if egg laying rates differed in response to amino acid drop-out diets, we grouped these diets into two categories: those that induce a “fast-decline” in egg laying (including arginine, isoleucine, leucine, lysine, phenylalanine, threonine, tryptophan drop-out diets, and No AA diets), and those that induce a “slow decline” in egg laying (including methionine, histidine, and valine drop-out diets). Because the shapes of these curves were notably non-linear, we fit them with either linear, second order polynomial, third order polynomial, or 3-paramenter logistic curves. The logistic curves significantly improved model fit to the data for the No AA when compared to all other models, indicated by a dramatic reduction in AIC and BIC values (Table S1).

We then determined whether the fast and slow decline diets significantly altered egg laying dynamics by asking whether fitting coefficients specific to each diet group significantly improved the fit of the logistic model when compared to a model with shared coefficients. We found that fitting the data with separate coefficients for fast and slow decline diets significantly improved the fit of our logistic regression (Table 1), suggesting that the histidine, methionine, and valine drop-out diets induce slower declines in egg production relative to all other drop-out diets and to the diet with no amino acids (Figure 2A). This in turn suggests that the decline in egg laying is not equally as sensitive to the absence of each amino acid in the diet.

**Table 1.** Comparison of egg laying dynamics coefficients between fast decline and slow decline diets. Diets were compared using a logistic model, 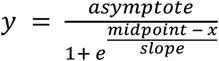, where y is the number of eggs laid per female and x is day.

|  | UPPER<br>ASYMPTOTE | MIDPOINT | SLOPE |
| --- | --- | --- | --- |
| **GROUPED** | 41.5 | -0.11 | -1.69 |
| **SLOW DECLINE DIETS** | 28.980 | 1.90 | -2.38 |
| **FAST DECLINE DIETS** | 21.57 | 1.49 | -0.65 |
|  | **F-value** | **df** | **p-value** |
| **GROUPED VS (SLOW DECLINE DIETS \| FAST DECLINE DIETS)** | 547.04 | 2 | **<0.001**<br>*** |

**Table 1:** Comparison of egg laying dynamics coefficients between White Dahomey and GCN2Δ flies. Diets were compared through a self-starting logistics model.

|  | WDAH |  |  | GCN2Δ |  |  |
| --- | --- | --- | --- | --- | --- | --- |
|  | Asymptote | Midpoint | Slope | Asymptote | Midpoint | Slope |
| GROUPED | 30.83 | 1.38 | -1.43 | 24.85 | 1.46 | -0.95 |
| SLOW DECLINE DIETS | 24.76 | 3.01 | -1.34 | 22.88 | 2.62 | -1.04 |
| FAST DECLINE DIETS | 24.14 | 1.56 | -0.72 | 21.12 | 1.38 | -0.51 |
|  | F-value | df | p-value | F-value | df | p-value |
| GROUPED VS (SLOW DECLINE DIETS \| FAST DECLINE DIETS) | 107.57 | 3 | <0.001 | 107.42 | 3 | <0.001 |

### TOR, but not GCN2, signalling mediates the difference in response to amino acid drop-out diets

Because amino acids are sensed within the cell by the GCN2 and TOR intracellular signalling pathways, we hypothesized that the difference in the rate of decline in egg laying between slow and fast-decline diets arises from differences in the sensitivity of either of these pathways to individual amino acids. To determine which of these two pathways was responses to the phenotypic differences in sensitivity to individual amino acids, we reduced either GCN2 or TOR signalling. We then compared egg laying when these females were fed on one of eight experimental diets: a diet containing all amino acids, a diet without amino acids, one of three diets that induced a fast decline in egg production (arginine, leucine, or phenylalanine drop-out diets), or one of three diets that induced a slow decline in egg production (histidine, methionine, or valine drop-out diets). The three fast-decline diets were chosen because they showed biochemical similarities with histidine, methionine, and valine: histidine and arginine are cationic amino acids, methionine and phenylalanine are hydrophobic amino acids, and valine and leucine are branched chain amino acids. If the differences in the rates of decline in egg production were due to differences in sensitivity of either GCN2 or TOR signalling to those amino acids, we would predict that reducing the signalling of these pathways would result in similar rates of decline across all amino acid drop out diets.

To alter GCN2 signalling, we compared egg production in females that carry a null mutation for GCN2 (GCN2Δ) to wild type females with the same genetic background, wDah. When comparing the response of egg laying across diets in this experiment, we found that the valine drop-out diet no longer induced a slow decline in egg laying for either genotype (Figure S1, F3 = 1.48, p value = 0.22). For this reason, we eliminated the valine and leucine drop-out diets from our analysis and from subsequent experiments.

To ensure that any differences in egg laying between genotypes did not result from differences in ovary size, we first dissected out the ovaries of GCN2Δand wDah females and counted the number of ovarioles, a commonly used measure of ovary size (Hodin & Riddiford, 2000; Sarikaya et al., 2012; Mendes et al., 2016). The number of ovarioles per ovary of GCN2Δ females did not differ from wDah females (Figure S2, χ^2^_1_ = 0.44, p value = 0.51). However, the number of eggs laid by GCN2Δ females on all diets decreased faster than the wDah controls (Figure 3A, Table 2), resulting in fewer eggs produced over the 8 days of assay (Figure 3B, Table S2). Even so, GCN2Δ females on the methionine and histidine diets still exhibited a slower rate of decline than those on the arginine and histidine drop-out diets (Figure 3A, Table 2). As a result, within the same genotype both the control and GCN2Δ flies laid significantly more eggs on the methionine and histidine drop-out diets than on the arginine and leucine drop out diets (Figure 3B). This suggests that GCN2 is required for normal rates of egg laying, but that it is not required to set the differences in rates of decline in egg laying among amino acid drop-out diets.

**Figure 3:**
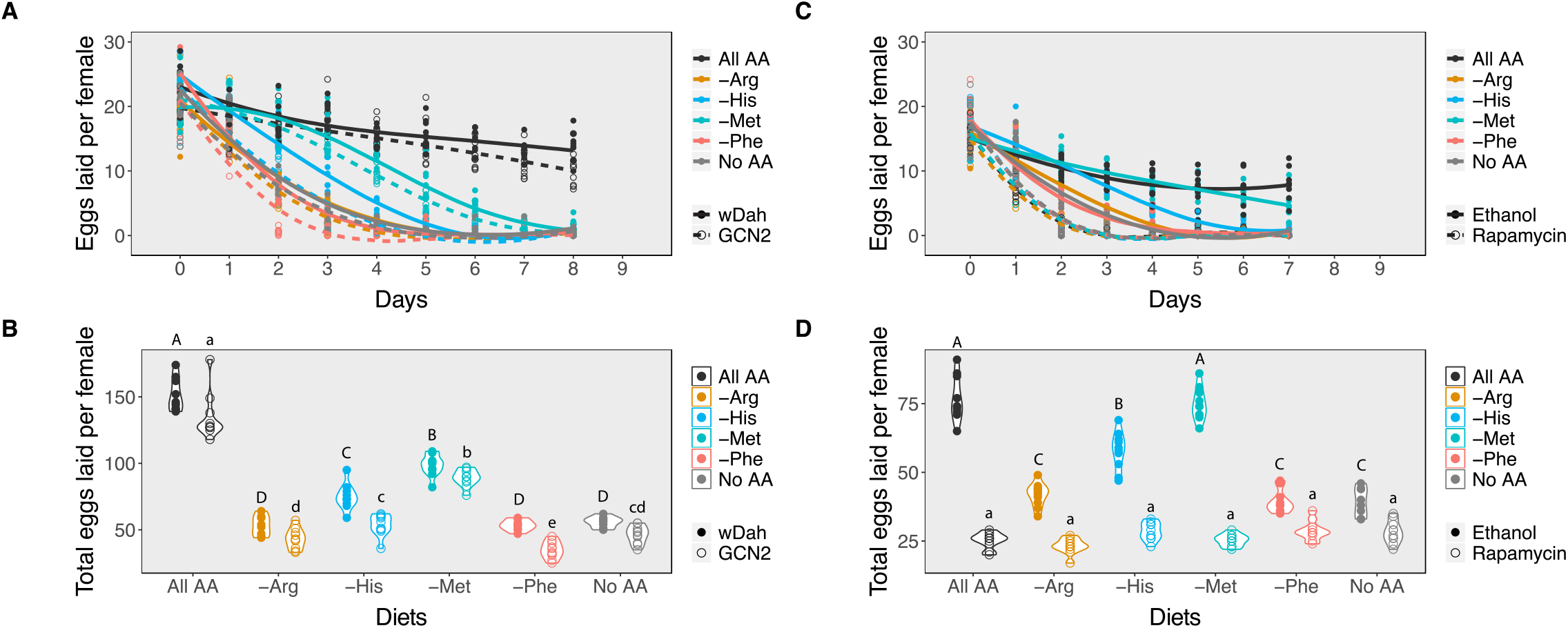
Inhibiting either the GCN2 or Target of Rapamycin pathways alters the impact of diet or egg laying. A) Changes in the rate of decline in egg laying in response to amino acid availability in the diet between control (wDah, solid lines) and GCN2Δ flies (dashed lines). B) Mean number of eggs laid per female in response to amino acid availability between control (wDah, closed circles) and GCN2Δ flies (open circles). C) Changes in the rate of decline in egg production in response to amino acid availability in the diet between ethanol-treated Red Dahomey flies (solid lines) and rapamycin-treated Red Dahomey flies (dashed lines). D) Mean number of eggs laid per female in response to amino acid availability between ethanol-treated Red Dahomey flies (open circles) and rapamycin-treated flies (closed circles). In B) and D), diets that have differing letters are significantly different within a genotype or treatment, as determined from post hoc pairwise comparisons derived from a generalised linear model assuming a Poisson distribution (Table S2). Capital letters indicate significance groups across diets for the control flies. Lower cases letters indicate significance groups for the GNC2 mutant (B) or rapamycin-treated (D) flies across diets.

**Table 2.** Comparison of egg laying dynamics coefficients between Ethanol and Rapamycin treated flies. Diets were compared through a self-starting logistics model.

|  | ETHANOL |  |  | RAPAMYCIN |  |  |
| --- | --- | --- | --- | --- | --- | --- |
|  | Asymptote | Midpoint | Slope | Asymptote | Midpoint | Slope |
| GROUPED | 28.49 | 0.55 | -1.72 | 17.35 | 1.03 | -0.30 |
| SLOW DECLINE DIETS | 19.34 | 2.74 | -2.00 | 16.82 | 1.06 | -0.29 |
| FAST DECLINE DIETS | 19.21 | 1.48 | -0.78 | 17.58 | 1.02 | -0.30 |
|  | F-value | df | p-value | F-value | df | p-value |
| GROUPED VS (SLOW DECLINE DIETS \| FAST DECLINE DIETS) | 128.25 | 3 | <0.001 | 2.37 | 3 | 0.07 |

Next, we assessed the effects of reducing TOR signalling on egg production on our six experimental diets. Since mutants for this pathway are usually developmentally lethal, we added rapamycin, a drug that inhibits this pathway, to the adult diets only. For our controls, we added the same volume of the carrier, ethanol, to the diets. We observed that when rapamycin was added to the diets, egg production declined at the same fast rate across all diets tested (Figure 3C, Table 3). Indeed, the decline in egg production in females treated with rapamycin was even faster than in females treated with ethanol on the no amino acid, arginine, and phenylalanine diets. This resulted in similarly low mean egg production per female across all diets (Figure 3D, Table S2), and suggests that the differences in decline in egg production across amino acid drop-out diets result from differences in the extent to which the absence of an amino acid can reduce TOR signalling.

### Interactions between cellular amino acid sensing pathways and egg chamber development

We can use the differences in the way GCN2 and TOR have been reported to affect egg chamber development to predict what we would expect to happen to the percentage of eggs in early, yolk-forming, and late-stages of egg development if the response to slow versus fast decline diets were due to TOR, but not GCN2, signalling. GCN2 signalling in the adult adipocytes represses both germline stem cell maintenance and ovulation (Armstrong et al., 2014, Figure 4A). Relative to control flies on diets containing all amino acids, we would predict that GCN2 null females could potentially exhibit a decrease in the percentage of egg chambers in the late stage of egg development on amino acid depleted diets (Figure 4B). This could occur since without GCN2, the stress response to uncharged tRNAs would not be induced. This would continue to trigger ovulation and deplete late egg chambers stages. TOR signalling is necessary for germline stem cell maintenance, for egg chamber growth, and for egg chambers to progress to the yolk-forming stages (Lafever et al 2010, Figure 4A). If TOR signalling differed in sensitivity to individual amino acids, we would predict that the percentage of egg chambers in yolk forming stages, and by association late stages, would decline more slowly in females offered the diets that induce a slow decline in egg laying when compared to females offered diets that induce a fast decline (Figure 4B). Rapamycin treatment would eliminate the differences between diets, and so, would result in an equal loss of yolk forming stages across all diets.

**Figure 4:**
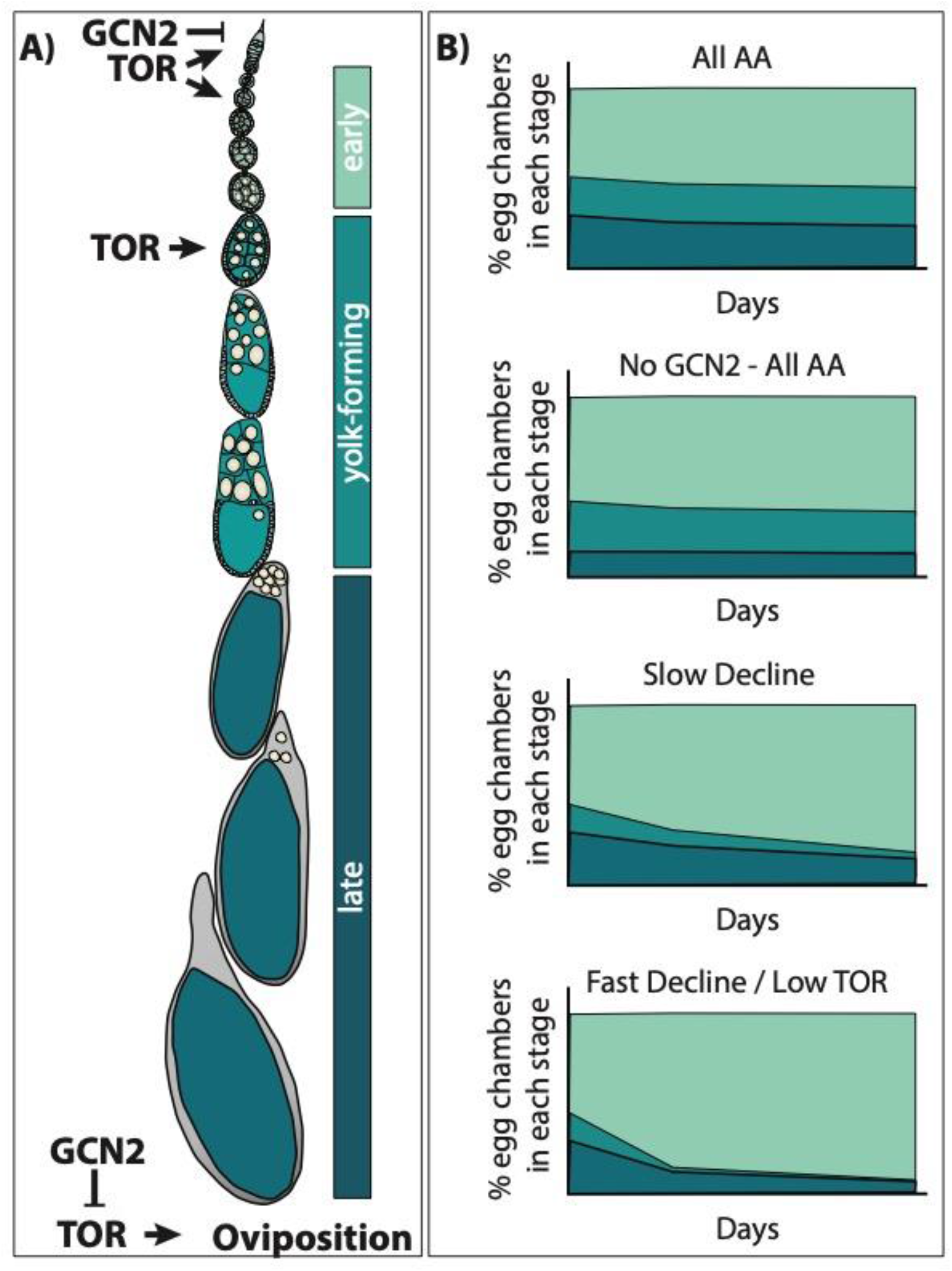
Predicting how diets that induce a slow versus fast decline impact egg laying via the TOR pathway. A) Egg chamber development can be broadly divided into three stages: 1) early stages (stages 1-7), yolk-forming stages (stages 8-10), and late stages (stages 11-14). TOR signalling promotes germline stem cell maintenance, egg chamber growth, and the transition into yolk-forming stages (Lafever et al 2010). Both TOR and GCN2 regulate oviposition; TOR signalling promotes oviposition while GCN2 signalling inhibits TOR to suppress oviposition (Armstrong et al 2014). B) GCN2 mutant females (No GCN2 – All AA) should have normal stem cell numbers and should not show repression of oviposition. Relative to wild type females on diets containing all amino acids (All AA), this might result in faster rates of oviposition, reducing the percentage of late stage egg chambers. We would expect ovaries from females on amino acid drop out diets (Slow and Fast Decline) to have reduced percentages of egg chambers in the yolk-forming and late stages relative to ovaries from females on All AA. If this difference is due to differences in the sensitivity of the TOR pathway across amino acids, then we would predict that on the slow decline diets we would see a more gradual decline in the percentage of yolk-forming egg chambers than on the diets that induce fast decline in egg laying. Treatment with rapamycin (low TOR) should reduce or eliminate the differences between slow and fast decline diets.

To test our predictions, we compared the developmental stages of egg chambers when females were subjected to one of six diets, all AA, no AA, or drop-out diets for arginine, phenylalanine, methionine, or histidine. We further reduced amino acid signalling either by using the GCN2Δ mutant flies or by adding rapamycin to the diet. Ovaries were imaged and egg chambers were then staged and allocated into one of the three groups, early, yolk-forming, and late stages.

In the ovaries of control white Dahomey females fed on a diet containing all amino acids, we found that 62% of the ovaries were in early stages, 22% were in yolk-forming stages, and 16% were in the late stages of egg chamber development (Figure 5A, Figure 6). These percentages remained relatively constant over the 7-day period examined. When these females were maintained on a diet that induced a slow decline in egg laying, either a diet lacking histidine or methionine, the percentage of yolk-forming and late-stage egg chambers declined with time (Figure 5B, Figure 6, Table 4). This resulted in a significant difference in the mean percentage of egg chambers in the late stages in comparison with ovaries from females fed a complete diet. Ovaries from females on the diets that induced a fast decline in egg laying, including diets lacking arginine, phenylalanine, or all amino acids, showed a similar rate of decrease in yolk-forming egg chambers, but a steeper decrease in late-stage egg chambers (Figure 5C, Figure 6, Table 4). The mean percentage of egg chambers in the yolk forming and late stages was significantly different to ovaries from females fed on diets with all amino acids or on diets that induced slow decline in egg laying. These data support our prediction that diets that induce a slow decline in egg production will also induce a slower decrease in the percentage of yolk-forming and late stage egg chambers than for flies on the amino acid dropouts that produce a rapid decline in egg laying (Figure 4B).

**Figure 5:**
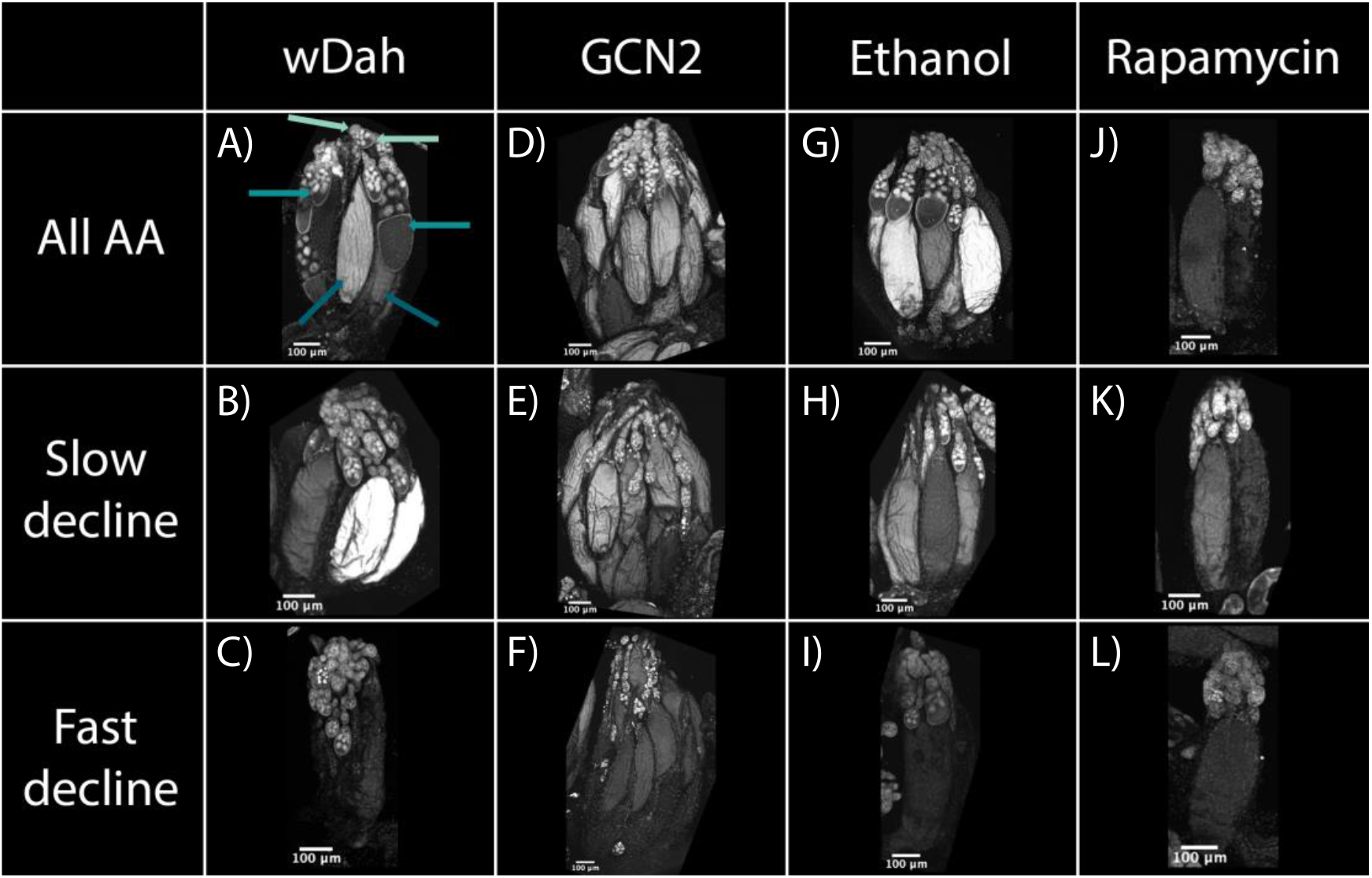
Egg chamber development changes with diet and with reductions in GCN2 and TOR signalling. Ovaries from white Dahomey (wDah, A-C), GNC2 mutant (D-F), red Dahomey fed diets laced with ethanol (Ethanol, G-I), and red Dahomey fed diets laced with rapamycin (Rapamycin, J-L) females. Females were fed either a diet consisting of all amino acids (All AA, A, D, G, and J), diets that induced a slow decline in egg laying (no histidine, B, E, H, and K), or diets that induced a fast decline in egg laying (no arginine, C, F, I, and L). All samples are stained with DAPI to visualise nuclei. In A), the light teal arrows point to early, the teal arrows point to yolk-forming, and the dark teal arrows point to late stage egg chambers. All pictures were taken after 7 days of diet exposure.

**Figure 6:**
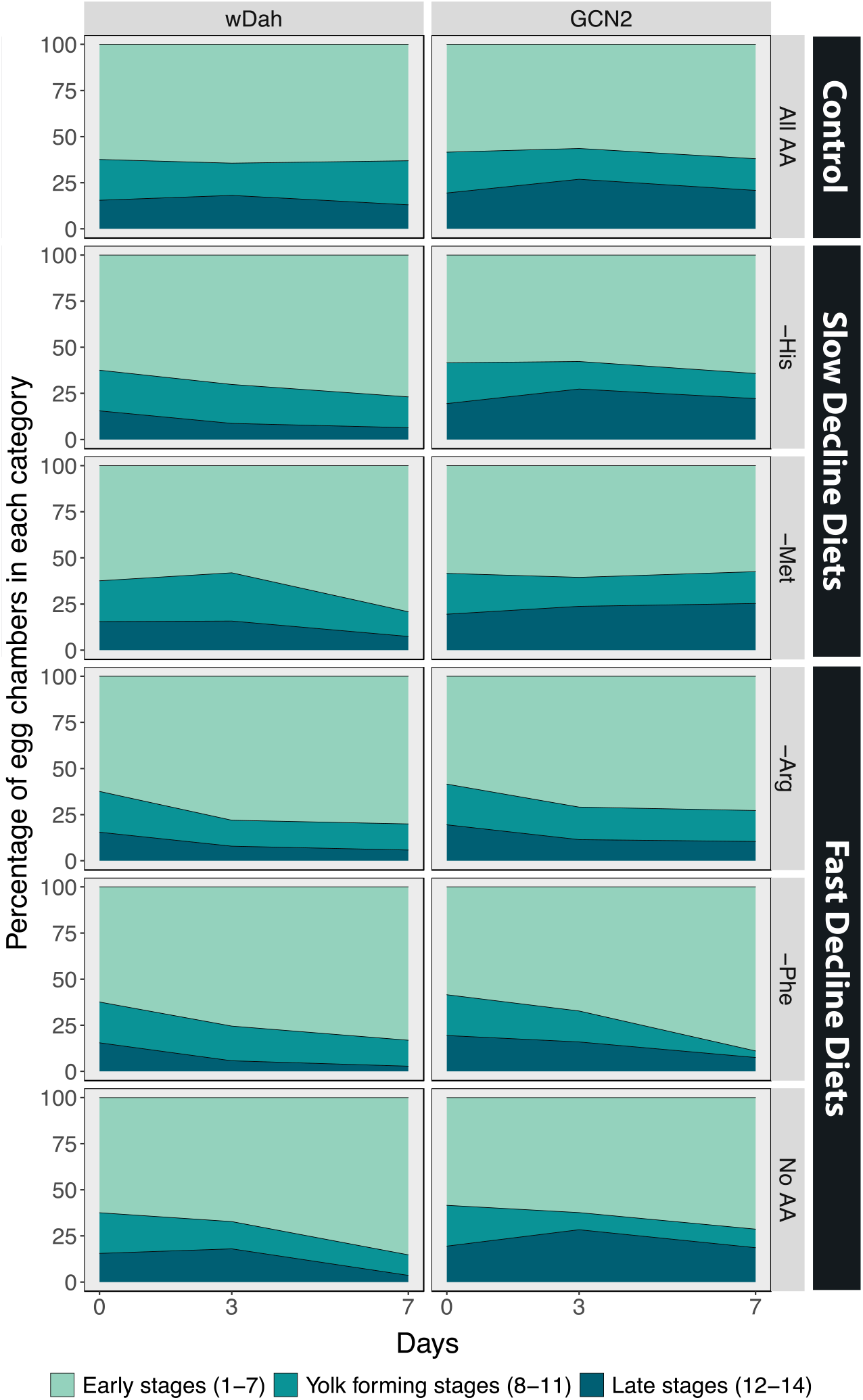
The percentage of egg chambers in each category is affected by diet, and these effects are partially mediated by GCN2 signalling. Comparison between the percentage of egg chambers between white Dahomey (wDah) and GCN2Δ flies.

**Table 4:** Drop-out diets that induce either a slow or fast decline (diet group) in egg laying differ in how they affect the percentage of egg chambers in each category over time in White Dahomey (wDah) control flies. Categories include early stages (egg chamber stage 1-7), yolk-forming stages (stages 8-11), and late stages (12-14). Diet groups that differ in the mean or slope for each category are indicated with different letters.

|  | wDah |  |  |  |  |  |
| --- | --- | --- | --- | --- | --- | --- |
|  | Chi-square | df | p-value |  |  |  |
| Day | 0 | 1 | 1 |  |  |  |
| Category | 9724 | 2 | <0.001 |  |  |  |
| Diet Group | 0 | 2 | 1 |  |  |  |
| Day*Category | 367 | 2 | <0.001 |  |  |  |
| Day*Diet Group | 0.3 | 2 | 0.74 |  |  |  |
| Category*Diet Group | 176 | 4 | <0.001 |  |  |  |
| Day*Category*Diet Group | 108 | 4 | <0.001 |  |  |  |
| Significance Groups Across Diet Groups |  |  |  |  |  |  |
|  | Early |  | Yolk-Forming |  | Late |  |
|  | Mean | Slope | Mean | Slope | Mean | Slope |
| All AA | A | A | A | A | A | A |
| Slow Decline Diets | B | B | A | B | B | B |
| Fast Decline Diets | C | B | B | B | C | C |

Egg chamber development in the ovaries of GCN2 mutant females differed from controls even on the diet that contained all amino acids. In these females, 57% of egg chambers were in early stages, 18% in yolk-forming stages, and 25% in late stages of development (Figure 5D, Figure 6). The increase in late stages of egg chamber development appear to reflect a block in ovulation (Figure 5D-F). Nevertheless, diets that induced fast decline still showed significant differences in the mean percentage and the rate of decline in both yolk-forming and late stages of egg chamber development from diets that induced a slow decline in egg laying (Figure 5E and F, Figure 6, Table 5). This suggests that while GCN2 does appear to increase the percentage of eggs in the late stages, it is not responsible for the differences in rates of decrease across amino acid drop-out diets.

**Table 5:**
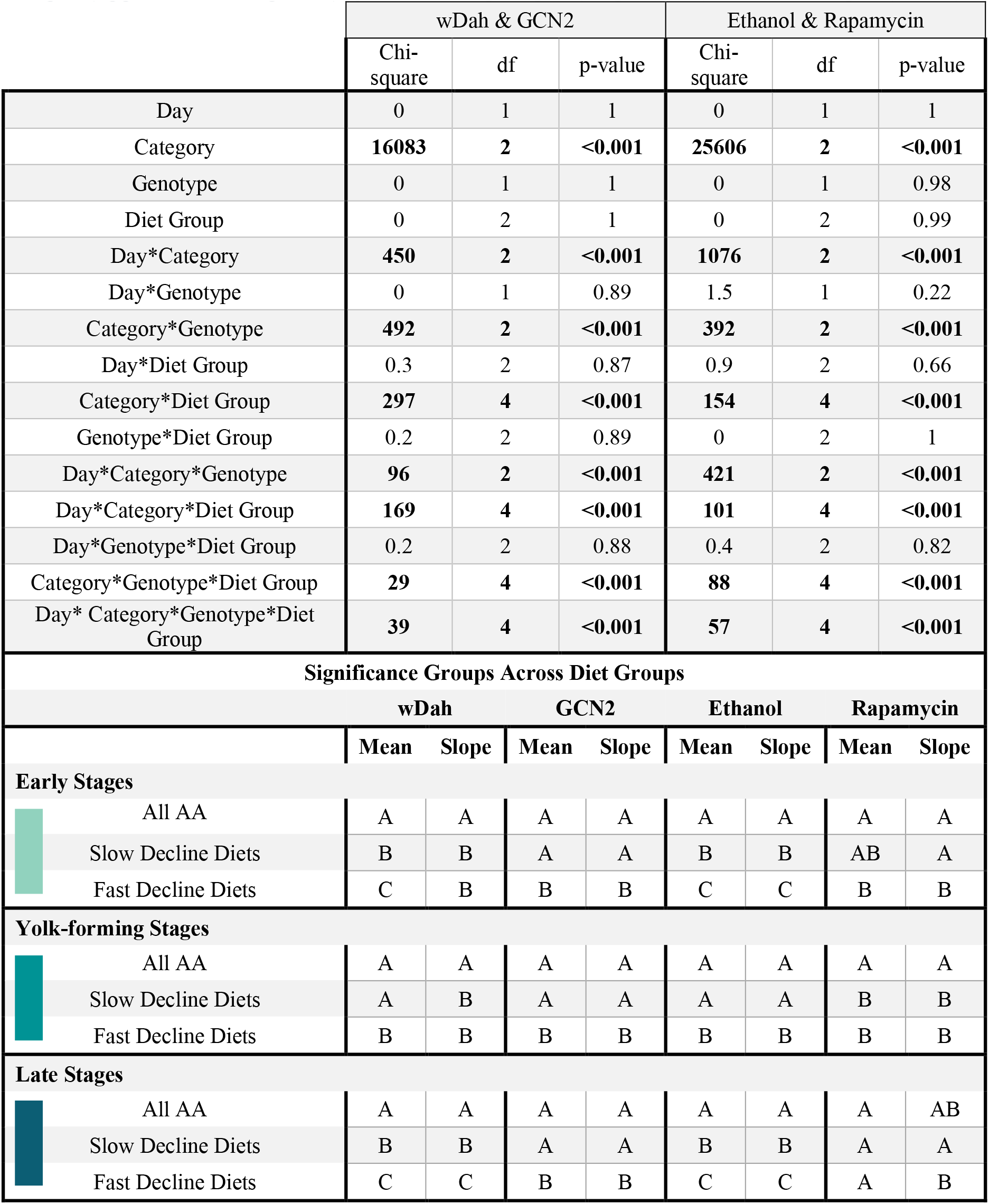
Manipulating both diet and either GCN2 or Target of Rapamycin signalling alters the percentage of egg chambers in each category over time. Categories include early stages (egg chamber stage 1-7), yolk-forming stages (stages 8-11), and late stages (12-14)

We next examined the effects of inhibiting TOR signalling on egg chamber development by adding rapamycin to the diet. As a control, we added the same volume of ethanol to the diets of red Dahomey females. We observed similar changes in the percentages of egg chambers in early, yolk-forming, or late stages when ethanol was added to the food of red Dahomey females as we did for the wDah females (GCN2Δ controls) (Figure 5G-I, Figure 7, Table 5).

**Figure 7:**
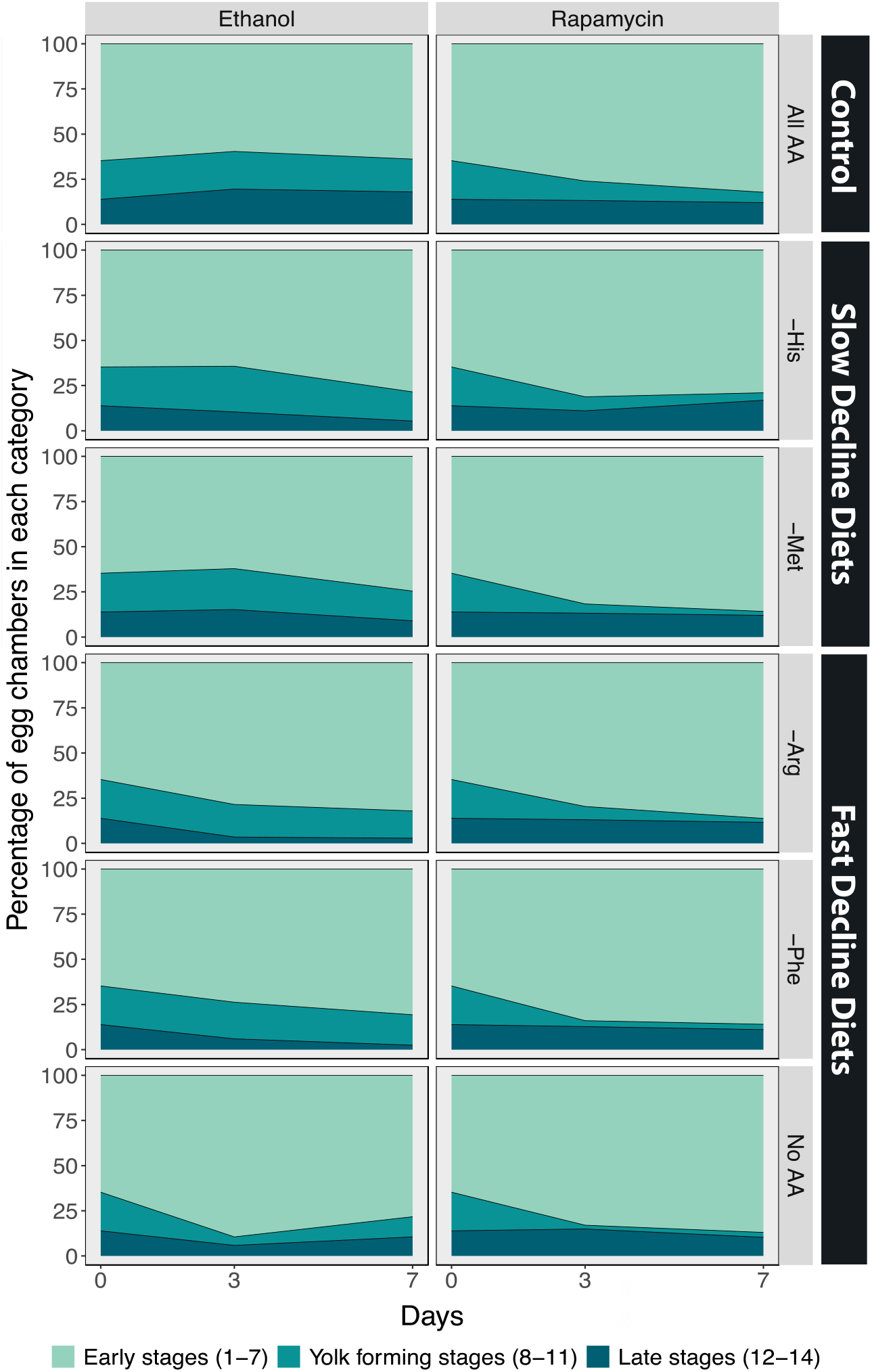
The percentage of egg chambers in each category differs among amino acid drop-out diets, and these effects are eliminated when Target of Rapamycin signalling is inhibited using the drug rapamycin. Comparison between the percentage of egg chambers between red Dahomey females treated with ethanol and treated with rapamycin (30μM).

Adding rapamycin caused rapid declines in the percentage of egg chambers in the yolk-forming stages in all diets (Figure 5J-L, Figure 7, Table 5). In addition, on diet without any amino acids the ovaries from rapamycin-treated flies had a higher percentage of late stage egg chambers (15%) relative to the ethanol treated controls (11%), indicative of a block in ovulation (Figure 5J-L, Figure 7). Finally, rapamycin treatment eliminated most of the differences in percentage of yolk-forming and late stage egg chambers between diets that induced a slow versus fast decline in egg laying (Figure 7, Table 5). Taken together, our data supports our predictions that the difference in decline in egg laying between slow and fast decline diets is mediated by TOR signalling. This provides further evidence that TOR signalling differs in its sensitivity across amino acids.

## Discussion

Accurately sensing the composition of the diet allows life-history traits to adjust with food quality and availability. How cellular signalling pathways can coordinate different sensitivities to amino acid availability and impact life history traits is still unknown. Here, we used the ovaries of *D. melanogaster* to understand how diet and the signalling pathways it activates impact a key life-history trait, fecundity. By manipulating individual essential amino acids of the diet, we found that egg laying shows amino acid-specific responses. Furthermore, we provide evidence that the TOR pathway is mediating difference in the response of egg laying to essential amino acids. These studies will provide insight into what limits the extent to which nutrient availability can be reliably conveyed to the organs of the body to regulate phenotypic responses.

Whether amino-acid sensing pathways are required at all for differences in sensitivity to the depletion of amino acids in whole animals is unclear. Previous studies have shown that removal of methionine from the culture medium of HEK293T cells blocks translation (Mazor et al., 2018). However, the block in translation does not appear to require either GCN2 or TOR pathways (Mazor et al., 2018). In the absence of methionine, GCN2 is not activated and TOR fails to inactivate. Instead, translation is blocked in these cells due to a lack of methionine-charged tRNAs necessary to initiate the process (Mazor et al., 2018). In addition, the absence of methionine results in an increase in DNA and histone methylation, which also inhibits translation. These studies suggest that at least in cell culture, the absence of methionine is not sensed. In a whole animal, this would mean that differences in phenotypic responses to methionine depletion would not be affected by altering either GCN2 or TOR signalling.

Our study suggests that these findings from cell culture studies do not apply to egg laying responses to amino acid depletion. While altering GCN2 signalling does not change the way egg laying responds to the depletion of single amino acids, we find evidence that the TOR pathway is important for the differences in response to methionine and histidine. This further suggests that TOR signalling itself might differ in sensitivity to individual amino acids, potentially driving a mismatch between the output of a nutrient sensing pathway and the true nutritional status of an organism.

*Drosophila* flies obtain amino acids primarily from the yeast that grows on the decomposing matter on which they live. Relative to the amino acid requirements encoded within the fly’s genome, methionine and histidine are the two most limiting amino acids in yeast (Piper et al., 2014, Gómez Ortega et al., 2021). Given that these two amino acids are present in lower abundance than the fly requires, potentially TOR signalling has adapted to be more tolerant of methionine or histidine deficiency via reduced sensitivity to these amino acids. The mechanisms that confer differences in sensitivity across amino acids are not yet understood.

TOR signalling acts on a number of cell types within the ovary to control egg chamber development (LaFever et al., 2010). In addition, TOR signalling acts in the fat body to regulate oviposition and is likely to also control the production of hormones, like the insulin-like peptides, ecdysone, and juvenile hormone, necessary for eggs to develop (Armstrong et al.,2014, Layalle et al., 2008, Hatem et al., 2015). Future studies manipulating TOR signalling in cell types known to be important for yolk-forming stages of egg development will highlight the cells responsible for differences in egg laying rates across diets.

While the GCN2 pathway did not drive the differences in egg laying across diets, flies mutant for GCN2 did show faster rates of decline in egg laying than wDah controls. These reduced egg laying rates appear to be caused by reduced oviposition frequencies in GCN2 mutants. This was surprising, as reducing GCN2 signalling in the fat body has previously been shown to relieve the impacts of amino acid depletion, which we anticipated would result in a reduction of egg production in the absence of essential amino acids (Armstrong et al., 2014). Our results suggest that GCN2 has a more complex role in egg laying than was previously thought. Whether this occurs via GCN2 activity in the fat body or in other cell types remains to be seen.

Finally, numerous studies have demonstrated cross talk between the TOR and GCN2 pathways (Rousakis et al., 2013; Yuan et al., 2017). Given this, it might seem surprising that they differ in the way they respond to the depletion of individual amino acids. However, TOR and GCN2 diverge in the pathways with which they interact. For example, TOR also responds to insulin signalling and carbohydrate levels (Jacinto and Hall, 2003). Moreover, null mutations in GCN2 are viable while null mutations in TOR are lethal. Future work will uncover the extent of overlap between these two pathways, and the contexts in which they diverge.

## Acknowledgements

We would like to thank members of the Mirth and Piper labs, past and present, for their help throughout this project. A special thank you to Amy Dedman and Tahlia Fulton for their unrelenting help in ovary collections. We would also like to thank Linda Partridge and Sebastian Grönke for sharing their unpublished GCN2 null stock. This research was funded by ARC (DP180103725) for CMS, ARC (FT170100259) for CKM, and ARC (FT150100237) and NHMRC (APP1182330) for MDP.

