## Supplementary Figures for "Target of Rapamycin drives unequal responses to essential amino acid depletion in egg laying"

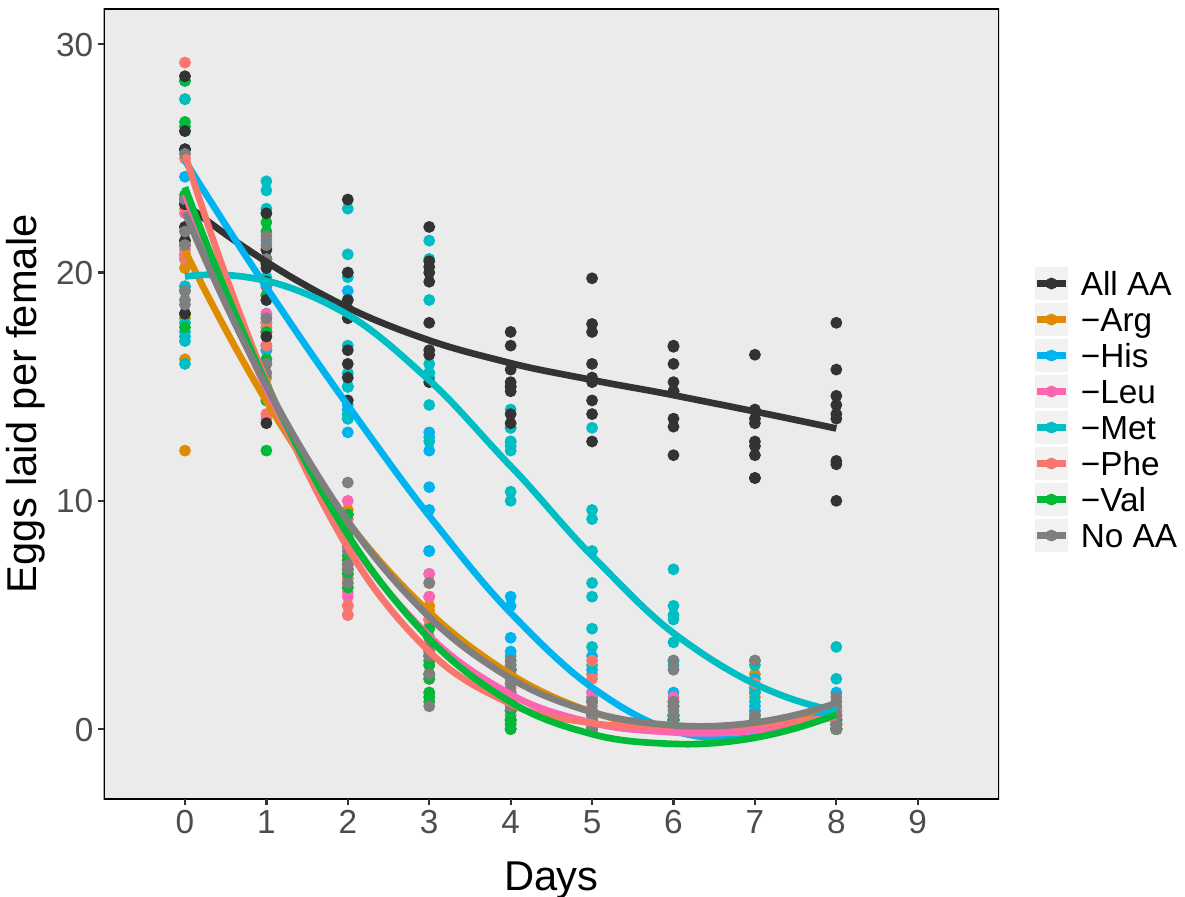
**Supplementary Figures & Tables**

**Figure S1: In the ovaries of white Dahomey females, a diet missing valine shows declines in egg laying with time that are indistinguishable with other diets that induce a fast decline in egg laying.**

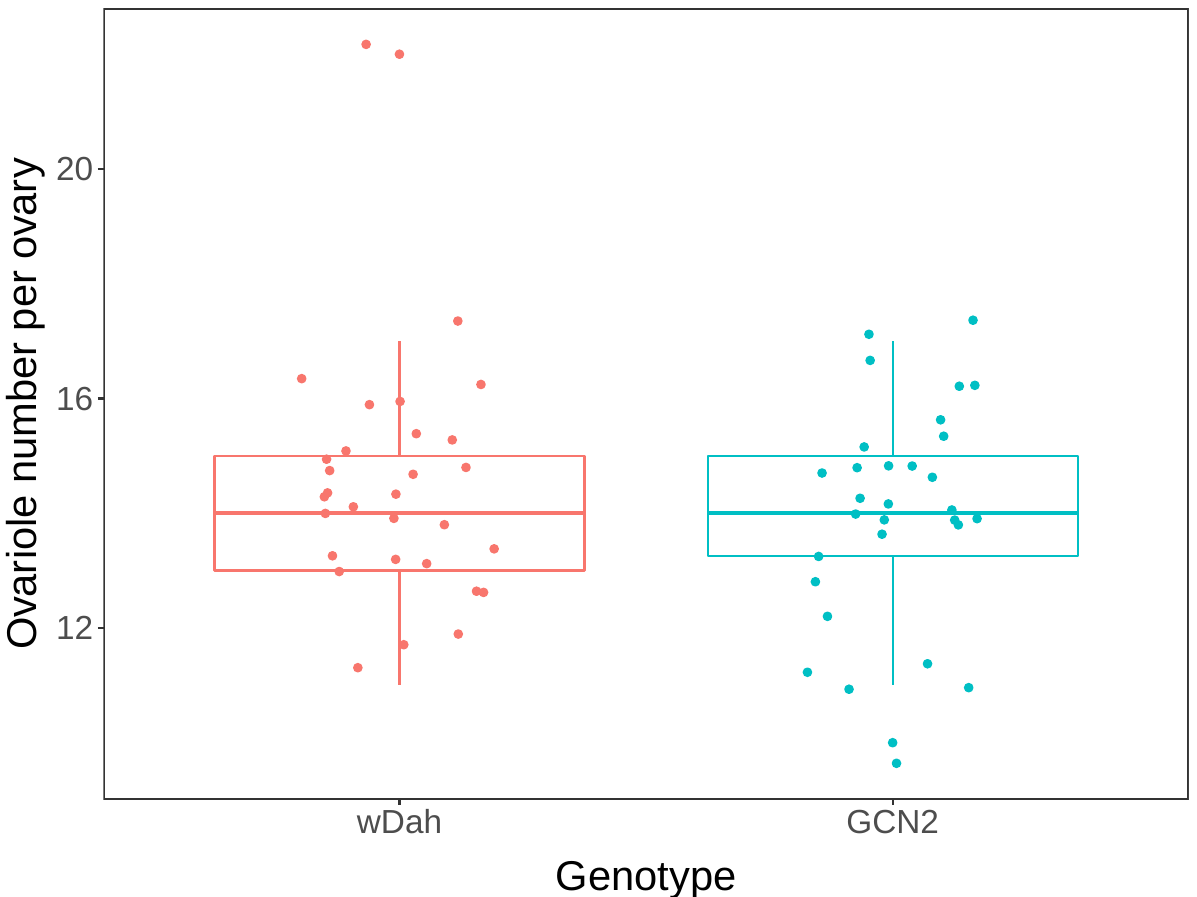

**Figure S2: Ovariole number does not differ between white Dahomey and GCN2 mutant females.**

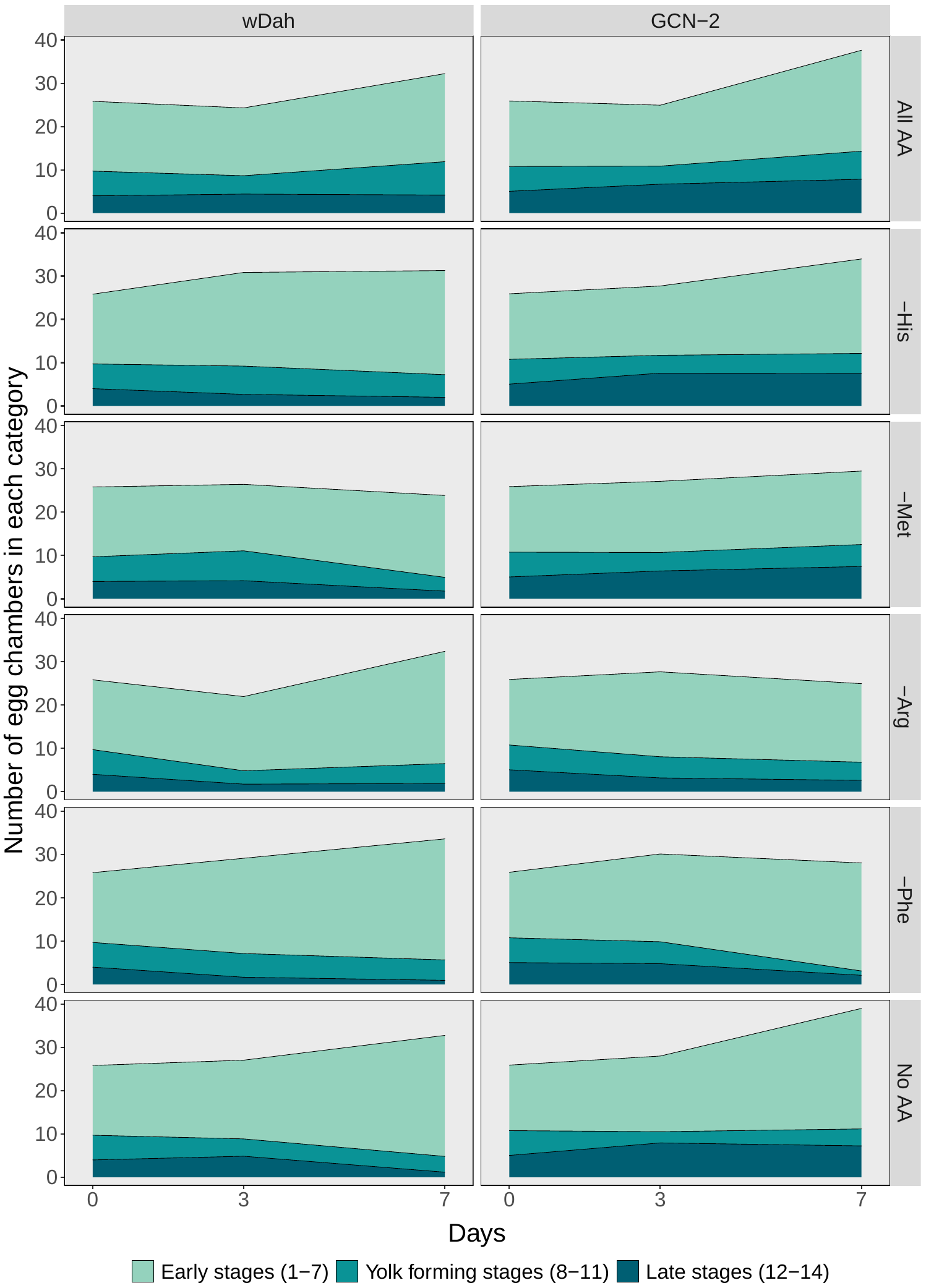

**Figure S3: The total number of egg chambers in each category over time in white Dahomey and GCN2 mutant females.**

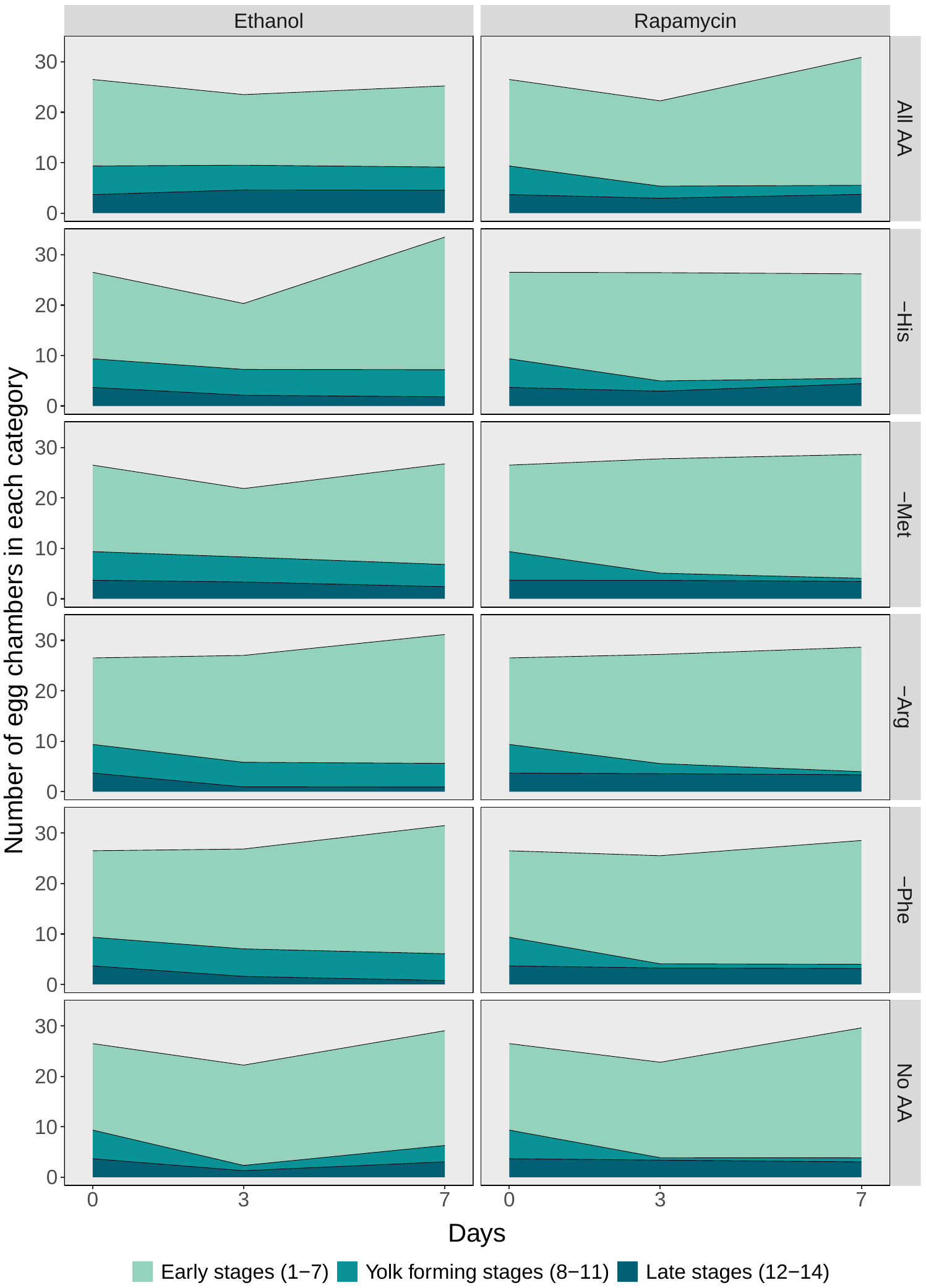

**Figure S4: The total number of egg chambers in each category over time in red Dahomey females treated with ethanol or rapamycin.**

Table S1 – Comparison of model fit for the No AA diet. Df: degrees of freedom, AIC: Akaike’s Information Criteria, BIC: Bayes Information Criteria

|  | df | AIC | BIC |
| --- | --- | --- | --- |
| Linear | 3 | 571 | 579 |
| Second Order Polynomial | 4 | 472 | 482 |
| Third Order Polynomial | 5 | 464 | 477 |
| Logistic | 4 | 408 | 419 |

|  | Total eggs laid | | |
| --- | --- | --- | --- |
|  | Chi-square | df | p-value |
| wDah & GCN2 | | | |
| Diet | **1669** | **5** | **<0.001** |
| Genotype | **76** | **1** | **<0.001** |
| Diet * Genotype | **25** | **5** | **<0.001** |
| Ethanol & Rapamycin | | | |
| Diet | **175** | **5** | **<0.001** |
| Genotype | **607** | **1** | **<0.001** |
| Diet * Genotype | **112** | **5** | **<0.001** |

Table S2: The total number of eggs laid across diets differs between wDah and GCN2 genotypes, and between ethanol and rapamycin treatments. Data are fit with generalised linear models assuming a Poisson distribution.
